# Rice Cultivars Carrying the Semi-Dwarfing Allele Enables High Yield without Lodging under Hairy Vetch–based Green Manure

**DOI:** 10.64898/2026.05.25.727576

**Authors:** Hirofumi Fukuda, Toshihiro Sakamoto, Akari Fukuda, Daisuke Ogawa

## Abstract

Green manure is a promising strategy for reducing dependence on chemical fertilizers in crop production. However, vigorous growth due to green manure often leads to high yield accompanied by lodging in rice, hindering its practical use in rice cultivation. Here, we show that rice cultivars carrying a semi-dwarfing *sd1/ga20ox2* allele achieve high grain yield without lodging under hairy vetch–based green manure conditions. The leading Japanese cultivar Koshihikari exhibited enhanced vegetative growth, increased panicle number, and consequently higher grain yield and quality under green manure conditions in 2023 and 2024 compared with chemical fertilizer management, although this was accompanied by increased culm length and widespread lodging. Among the four *GA20-oxidase* genes, green manure significantly upregulated *Sd1*/*GA20ox2* mRNA levels. A temperate *japonica* cultivar, Nijinokirameki, and an *indica* cultivar, Hokuriku-193, carrying a non-functional *sd1*/*ga20ox2* allele exhibited no lodging under hairy vetch–based green manure management while achieving improved yield performance. Notably, yields obtained under our hairy vetch–based cultivation system were comparable to or exceeded a recently reported high-yield benchmark observed for Hokuriku-193 under chemical fertilizer management in the same region of Japan. These findings suggest that cultivars harboring non-functional *sd1*/*ga20ox2* alleles enable the practical implementation of annual hairy-vetch-rice rotation for sustainable rice production.

## Background

Rice (*Oryza sativa* L.) is a major staple crop all over the world, and the rising global population demands a consistent increase in its production to ensure food security (Van Nguyen and Ferrero, 2006; Muthayya et al., 2014). Over the 25-year period from 2010 to 2035, rice demand in Africa is predicted to increase by 30 Mt while Asian rice intake is forecasted to rise by 77 Mt, accounting for 67% of the global increase of demand (IRRI 2010). In addition to the need for increased food supply, especially in regions where rice serves as the primary calorie source (Gnanamanickam, 2009), improving rice yield is important for farmers who depend on income from its sale. Thus, improved yields can alleviate hunger, improve farmer incomes, and contribute to economic growth.

Green manure (e.g., leguminous plant material) offers a promising solution to the challenge of improving rice yields (Islam et al., 2019). It enhances soil organic matter, increases nutrient availability, and improves soil properties (Kumar et al., 2014; Xie et al., 2016; Luo et al., 2020; Lei et al., 2022). Green manure is an eco-friendly alternative to chemical fertilizers because it biologically fixes nitrogen (N), making it a potentially renewable N source in an on-farm production system (Pakeerathan et al., 2009; Raheem et al., 2019). Cherr et al. (2006) noted that the use of animal manure also can meet soil N fertility demands but may result in soil phosphorus (P) loading owing to the imbalance between its low N/P ratios and the P requirements of crops (Royer et al., 2003; Hao et al., 2004); in addition, animal manure may cause soil salinization because of the resulting high ion concentrations (Hao and Chang, 2003).

Despite the benefits of green manure, the molecular basis for its effects on yield and grain appearance quality remains unclear. Furthermore, its use poses a challenge. For example, excessive crop growth can lead to lodging (the bending or falling over rice plants), which decreases both yield and harvest efficiency (Watanabe, 1984; Shah et al., 2017). One effective approach to addressing the lodging problem is to make culm length and plant height shorter through the use of semi-dwarf alleles (Sasaki et al., 2002; Murai et al., 2004; Tomita, 2009; Liu et al., 2018). For example, the *semi-dwarf 1* (*sd1*) allele, which harbors a deletion (∼380 bp) in the first exon of *gibberellin 20 oxidase-2* (*GA20ox2*), enhances lodging resistance in rice even under a high-N-fertilizer condition (reviewed by Nagai et al., 2018). Thus, taking advantage of *Sd1*/*GA20ox2* variants may play a crucial role in alleviating lodging risk under green manure treatment, facilitating its use in sustainable rice farming systems. In this study, we first examined the overall effects of green manure (hairy vetch) treatment on plant growth, yield, and lodging resistance in the leading Japanese rice cultivar Koshihikari. This cultivar contains a non-dwarf/functional allele of *Sd1/GA20ox2*. Additionally, we explored the impact of green manure treatment on leaf transcriptome profile at vegetative and panicle formation stages, N content indicators at the panicle formation stage, grain protein content, and grain appearance quality. Subsequently, we extended the investigation of lodging and yield performance to other high-yielding *japonica* and *indica* rice cultivars that possess a semi-dwarf (non-functional) allele of *sd1*/*ga20ox2*.

## Materials and methods

### Cultivation and decomposition conditions of hairy vetch plants grown under experimental field conditions

Hairy vetch (*Vicia villosa*), a legume species, was grown as a cover crop in paddy fields in Kannondai, Tsukuba City, Ibaraki Prefecture, Japan (36°01′29″N, 140°06′25″E, 21 m a.s.l.). Nutrient element contents in the field soils were analyzed by Tokachi Federation of Agricultural Cooperatives (https://www.nokyoren.or.jp/; Hokkaido, Japan), described in Supplementary Table 1. The seeds (30 kg/ha; Fujiemon [Massa], Snow Brand Seed Co., Ltd.) were sown in field by hand on October 21, 2022, for rice cultivation in 2023 and on October 22, 2024, for that in 2025. For rice cultivation in 2024, we sowed that same amount of hairy vetch seeds on October 31 and again on November 15, 2023 (i.e., 60 kg/ha in total) since the germination rate of the first-sown seeds was low. Based on visual observation in the field, the biomass produced by this first sowing was negligible, and thus only a very small amount of biomass from the first sowing was returned to the soil. The plants were grown for 5 months from October to April in 2023, 2024, and 2025 and then directly plowed into the soil in the Kannondai paddy fields. Shoot biomass for hairy vetch was estimated using the dry weights of the samples analyzed for nitrogen content. Each sample was collected from a 25 cm × 25 cm area, as one biological replicate, just before the remaining hairy vetch plants were plowed into the soil. The dry weight of these samples was then used to extrapolate shoot biomass in tons per hectare.

### Rice cultivation conditions under chemical fertilizer use and green manure treatment under experimental field conditions

Seeds of three rice cultivars [the Japanese leading cultivar Koshihikari, a high-yielding temperate *japonica* cultivar Nijinokirameki (Ishimaru et al., 2023), and a high-yielding *indica* cultivar Hokuriku 193 (Goto et al., 2009)] were soaked in water at 30 °C for 2 d, sown in trays filled with synthetic granular soil (Sumitomo Chemical Company Co., Ltd., Tokyo, Japan), and incubated at 30 °C in the dark for 2 d. The seeds were sown on April 17 or 24, 2023, on April 5 or 22, 2024, and on April 28, 2025. Those trays placed in a field for nearly 3 weeks to 1 month and then transplanted (18 cm apart × 30 cm apart, planting density 18.5 hills/m^2^) into the desired field conditions and grown for approximately 5 more months from mid-May to September. A normal paddy field condition (fertilization with 56:176:56 kg/ha of N:PO_4_:K) was set as the control condition. No chemical fertilizers were added to the green manure–treated soil. Meteorological data of daily average temperature and relative humidity at 1-d intervals were obtained from the Japan Meteorological Agency during cultivation seasons (Supplementary Fig. 1).

### Manual assessment of agronomic traits of rice plants

Agronomic traits were evaluated as described in previous studies (Ogawa et al., 2021; Fukuda et al., 2025). Generally, one biological replicate was based on one bulk of three seedlings. Days to heading was defined as the number of days from sowing to the emergence of the first panicle in more than half of the plants in each cultivar. Culm length and panicle length of the longest culm per plant were measured using a ruler, and tiller (panicle) number was counted from 1/2 to 2 months after heading. Dry weight was measured after drying mature shoots (i.e., aboveground plant parts) for at least 1 month. Grain weight was measured after threshing the plant bulks and divided by the number of plants per bulk to calculate the grain weight per plant. Estimated grain yield (t/ha) was calculated using the grain weight per plant bulk (with the average bulk assumed to cover 5.4 × 10^−2^ m^2^ of field area). Grain appearance quality was assessed using 96-dpi scanned images of over 400 seeds per plant as one biological replicate. Seeds were collected from panicle bulks, and the percentage of perfect grains was calculated using a grain discriminator (RGQI 100B, Satake Corp., Hiroshima, Japan). Perfect grains were defined as non-chalky grains having a normal shape, as in our previous study (Fukuda et al., 2025). Chalky grains were classified into four types: white-milky, basal-white, white-belly, and white-back. Grain length, grain width, and grain thickness were also measured using the same grain discriminator.

### Measurement of nitrogen

Samples for N analysis were prepared from whole shoots of 5-month-old hairy vetch plants and 3-month-old rice plants grown in the experimental fields. Sample processing and N quantification followed Arai-Sanoh et al. (2014), with slight modifications to the sample processing method. Briefly, shoots were oven-dried at 30 °C for more than 2 weeks, weighed, and ground to fine powder using a mill (CMT TI-100; Cosmic Mechanical Technology, Fukushima, Japan). N concentration on a dry-weight basis was then determined using a Sumigraph NCH-22 analyzer (Sumika Chemical Analysis Service, Ltd., Tokyo, Japan), using the same method as in Arai-Sanoh et al. (2014).

### Aerial image capturing

SOTEN (ACSL, Tokyo, Japan) camera-equipped unmanned aerial vehicles (UAVs) were used to obtain aerial multispectral images for calculating chlorophyll index–green (CI_green_), and Phantom 4 Pro V2 (P4V2; DJI, Shenzhen, China) was used to obtain aerial color images for measuring canopy height using Structure-from-Motion/Multi-View-Stereo (SfM-MVS) analysis. The multispectral camera unit mounted on SOTEN units (CX-GB300; Xacti, Osaka, Japan) was equipped with a tri-band bandpass filter (TB550/660/850; Midwest Optical Systems, Palatine, Illinois, USA) that captures spectral images of green (543–558 nm), red (653–668 nm), and near-infrared (NIR; 835–865 nm) with a 1-inch imaging sensor (image resolution 2720 × 1814 pixels). The SOTEN was manually operated to capture 222 spectral images at an altitude of 30.5 m (ground resolution: 1.37 cm/pixel) on June 23, 2023. The onboard camera of the P4V2 also featured a 1-inch imaging sensor (image resolution 5472 × 3648 pixels). The P4V2 was manually operated to capture 94 color images at an altitude of 33.8 m (ground resolution: 0.9 cm/pixel) on September 4, 2024, for observing the canopy height of rice crops during the ripening stage. The P4V2 was automatically controlled by dedicated flight control software, DJI GS Pro (DJI), to capture 134 color images at an altitude of 32.2 m (ground resolution: 0.8 cm/pixel) on May 7, 2025, after the hairy vetch plants had been plowed under green manure–treated condition but before rice planting, for measuring the elevation of the ground surface after leveling.

### Processing of captured aerial images

The orthomosaic-processed spectral reflectance image and digital surface model (DSM), which represent three-dimensional height information of the subject, were generated from overlapping aerial images through SfM-MVS analysis using Agisoft MetaShape Professional v.2.2.0 software (Agisoft, St. Petersburg, Russia). As for SOTEN data, spectral images were converted to reflectance using light sensor and standard reflectance panel image information, and then made orthomosaic images with a modified workflow (Sakamoto et al., 2022). The following steps were (i) photo alignment, (ii) reflectance calibration using sun sensor data and standard reflectance panel images, (iii) DSM generation from tie points, and (iv) orthomosaic reflectance image generation. Since ground control point (GCP) information was not obtained during this flight, we manually selected tie points in ENVI v6.1 software (NV5 Geospatial, Broomfield, Colorado, USA) between reflectance images and performed geometric correction using a 1^st^-degree polynomial with the nearest-neighbor method to correct the pixel position of the green reflectance image relative to the NIR reflectance image. The procedures for creating the crop surface model (CSM) basically followed those described previously (Gitelson et al., 2003, 2005; Taniguchi et al., 2024). The DSM creation from high-resolution color images followed these sequential steps: (i) photo alignment, (ii) GCP input, (iii) point cloud generation, and (iv) DSM generation. The CSM was then created by taking the difference between the DSMs from September 4, 2024, and May 7, 2025, using ENVI.

### Quantification of CI_green_ and plant height

CI_green_ was calculated as the ratio of NIR to green reflectance values (Gitelson et al., 2003; Gitelson et al., 2005). Four replicated sampling areas (54 cm × 90 cm on the ground, each including six plants) were set up for each cultivar. The sampling areas were extracted from the CI_green_ image using ENVI software. Then, the mean CI_green_ was calculated for each single cut-out image. Plant height was measured using CSM data by setting a 44-m transect line across the field. The mean CSM value for each 10-cm grid along the transect line (*n* = 440) was calculated. Each cut-out CI_green_ image and grid sampling of CSM data was treated as a biological replicate for CI_green_ and plant height, respectively.

### Measurement of protein content in grains

Approximately 1000 grains, as one biological replicate, were measured by an NIR grain tester (AN-820; Kett Electric Laboratory Co., Ltd., Tokyo, Japan), following the manufacturer’s protocol. Rice grains provided by Kett and containing 0.63% protein, 13.5% moisture, and 17.6% amylose were used as a standard.

### SPAD measurement

SPAD values for the first fully expanded leaf from the top of the plant were recorded at the panicle formation stage. SPAD was measured at eight evenly spaced points on the flag leaf lamina at ∼1/3 of the blade length from the tip, avoiding the midrib. One plant per cultivar was randomly selected as a biological replicate; the mean SPAD of the eight points on the flag leaf was used as the value for that plant.

### RNA extraction

The largest leaves of vegetative Koshihikari plants at 47 days after transplanting (DAT) in 2023 and 38 DAT in 2024; and flag leaves at 5 days after flowering in 2024 (Supplementary Table 2) were used for the following processes. Each leaf sample was collected from 9:00 to 10:00 am as one biological replicate, immediately frozen in liquid N_2_, and stored at −80 °C until sample crushing. Frozen samples were crushed by a multi-specimen cell disruption device (BMS-A20TP; BMS, Tokyo, Japan) just before RNA extraction. Total RNA from each sample was extracted using a RNeasy Plant Mini Kit (Qiagen, Hilden, Germany) according to the manufacturer’s protocol.

### RNA-seq analysis

RNA-sequencing (RNA-seq) libraries were prepared using the NEBNext Ultra II Directional mRNA-seq kit for Illumina (New England Biolabs, Ipswich, MA, USA), following the manufacturer’s protocols. The libraries were sequenced using the Illumina NovaSeq X Plus. Sequencing was performed on the 25B flow cell with paired-end 150-bp and unique dual-index reads.

The reads were preprocessed by CLC Genomics Workbench version 25.0 (Qiagen). Reads were first trimmed using the following parameters: quality limit of 0.05, maximum number of ambiguities 2. Next, raw reads were mapped to the genome assembly and gene sets of Os-Nipponbare-IRGSP-1.0 with the following parameters: mismatch cost of 2, insertion–deletion cost of 3, length fraction of 0.8, similarity fraction of 0.8, and maximum number of hits of a read of 10. Then, uniquely mapped read counts were quantified. Genes with at least one raw read count in all of the library samples for each comparison were analyzed. Log_2_(fold change [FC]) and adjusted *P* values (Wald test using the “DESeq” function) were calculated from raw read counts by the R package “DESeq2” (version 1.42.1) in R software (version 4.3.2). Differentially expressed genes (DEGs) were extracted based on |log_2_(FC)| > 1 and adjusted *P* values (padj) < 0.05 compared to the control group.

Principal component analysis was performed using the count data transformed to the log_2_ scale, normalizing with respect to library size, by the rlog function in DESeq2. The first and second principal components (PC1 and PC2) were visualized using the plotPCA function in DESeq2.

## Enrichment gene ontology analysis

Gene ontology (GO) analysis of DEGs was conducted using ShinyGO v0.81 (Ge et al., 2020) with ‘osativa_eg_gene’ as the database ID and ‘Oryza sativa Japonica Group genes IRGSP-1.0’ for the genome assembly. The most significantly enriched GO terms were selected based on ‘FDR, sorted by Fold Enrichment’.

### Genotyping through DNA markers linked to a non-functional *sd1/ga20ox2* allele

To genotype the non-functional *sd1*/*ga20ox2* allele most often used in the Japanese breeding programs (a deletion-type mutant originating from the Taiwanese cultivar Dee-geo-woo-gen; Murai et al., 2004), a previously established forward primer (Sha et al., 2022) and a newly designed reverse primer (Supplementary Table 3) were used. The DNA fragments were separately amplified by PCR. The genomic region covering the deletion-type *sd1*/*ga20ox2* allele was amplified using PrimeSTAR HS DNA Polymerase (Takara Bio), according to the manufacturer’s protocol.

### Statistical analysis

All experimental data were averages from at least three independent experiments, and the values were subjected to statistical analysis through Student’s *t*-test, or the Tukey–Kramer test if ANOVA indicated a significant difference (*P* < 0.05) among group means overall.

## Results

### Growth and nitrogen concentrations of hairy vetch cultivated under field conditions

To evaluate the potential of hairy vetch as a green manure for rice cultivation, hairy vetch was cultivated in fields designated for subsequent rice planting (Fig. 1). Hairy vetch roots developed nodules, suggesting symbiotic rhizobia interaction that would facilitate atmospheric N fixation (Mylona et al., 1995; Zahran, 1999). Estimated shoot biomass (dry weight), N concentrations, and estimated N input (total N amounts) of hairy vetch plants just before being plowed were shown in Table 1. Each of values was 6.8±0.9 (t/ha), 0.038±0.003 (g/g dry weight), and 262.2±55.8 (kg/ha).

**Fig. 1.**
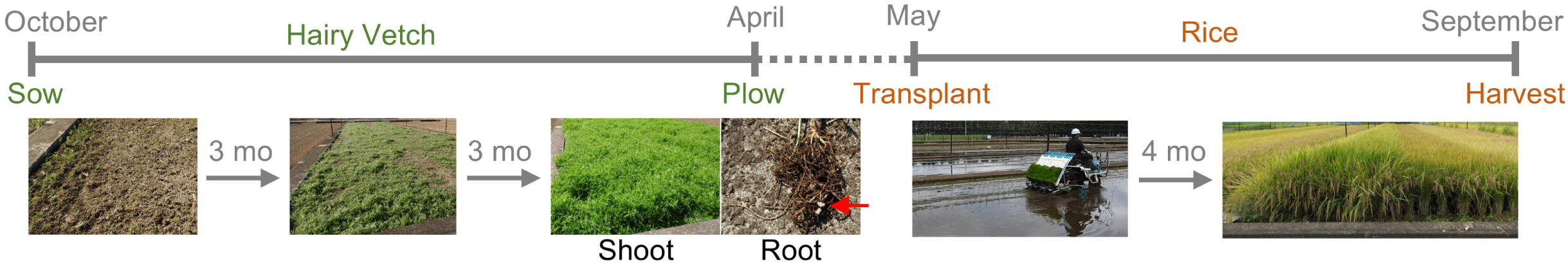
Scheme of cultivation processes from hairy vetch cultivation to rice plant harvesting. Red arrow indicates root nodules.

**Table 1.** Estimated shoot biomass, nitrogen concentration and estimated nitrogen input of hairy vetch plants. Hairy vetch plants were sampled just before the remaining plants were plowed into the soil. Mean values ± SD (*n* = 4) are shown.

| Estimated shoot<br>dry weight<br>(t/ha) | N conc.<br>(g/g DW) | Estimated N input<br>(kg/ha) |
| --- | --- | --- |
| 6.8 ± 0.9 | 0.038 ± 0.003 | 262.2 ± 55.8 |

### Effects of hairy vetch treatment on vegetative growth, yields, and grain appearance quality of Koshihikari under field conditions

In Koshihikari rice plants measured 46 DAT, the green manure treatment significantly increased CI_green_ compared to the use of chemical fertilizer, indicating higher chlorophyll levels (Fig. 2a). Aboveground weight, panicle number, and grain weight per plant were significantly increased under green manure treatment relative to chemical fertilizer treatment (Fig. 2b, c). Panicle length was not significantly changed by the green manure treatment. These findings suggested that the green manure treatment increased biomass and panicle number, leading to higher grain yield than under the chemical fertilizer treatment. We next assessed grain appearance traits between chemical fertilizer treatment and green manure treatment. The frequencies of chalky grains (e.g., those with white-belly and back) were significantly lower under green manure treatment, associated with more percentage of perfect grains and protein content in the grains (Fig. 2d–f). Furthermore, grain shapes and sizes including length, width, and thickness were comparable between the two treatments (Supplementary Table 4).

**Fig. 2.**
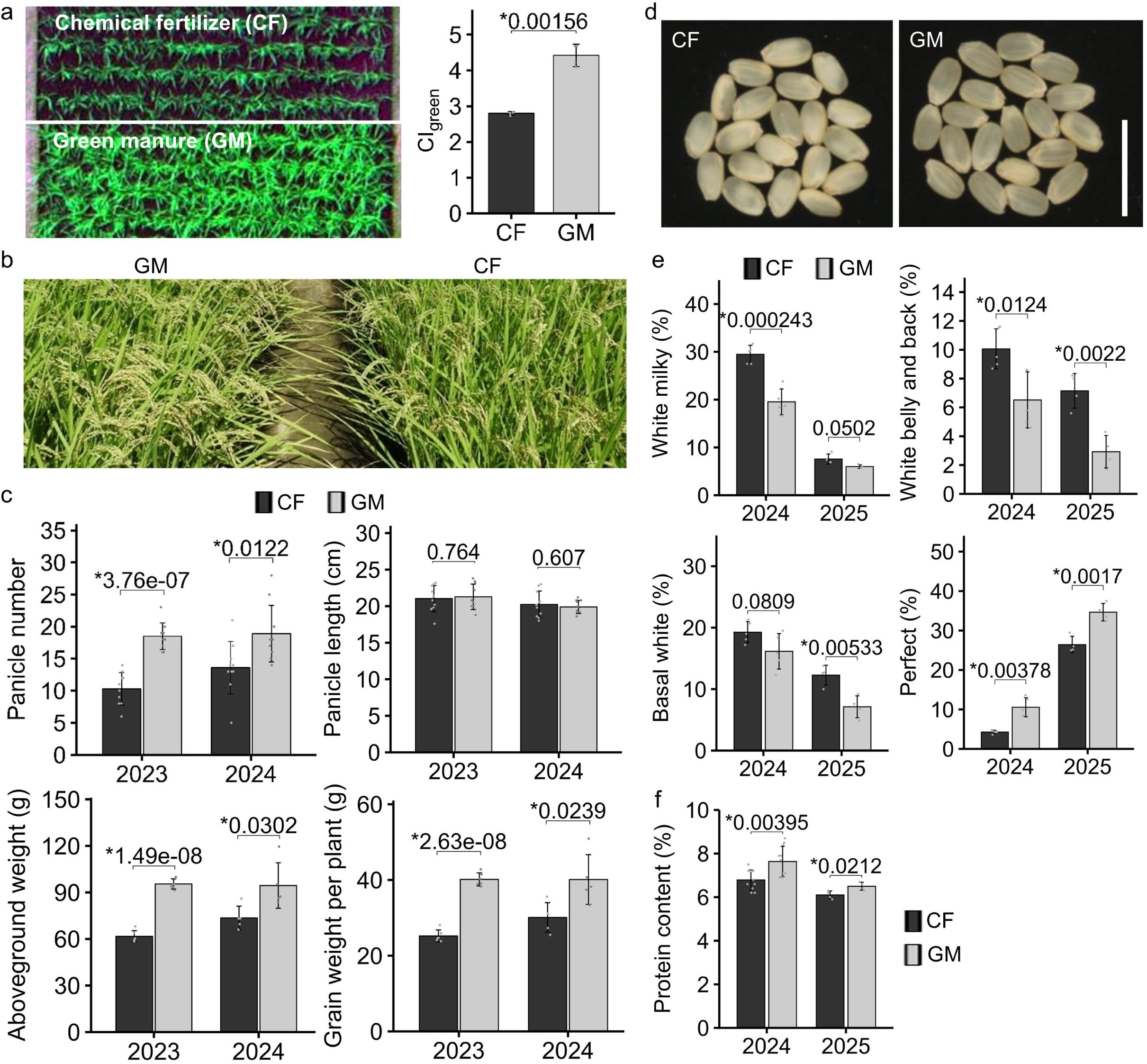
Agronomic traits related to yield and gran appearance quality in Koshihikari. (a) Vegetative growth under chemical fertilization (CF) and green manure treatment (GM) at 46 days after transplanting. Spectral reflectance image [R:G:B = ρ(Red):ρ(NIR):ρ(Green)] obtained by SOTEN and green chlorophyll index (CI_green_) of rice plants. Bars show mean values ± SD (*n* = 4). (b) Shoot growth at 94 days after transplanting in 2023. (c) Panicle number, panicle length, aboveground weight, and grain weight per plant at harvesting stage in 2023 and 2024. Bar plots indicate mean values ± SD (panicle number and panicle length, *n* = 10; aboveground weight and grain weight per plant, *n* = 6). (d) Grains of rice plants cultivated under experimental field conditions. Grains of plants cultivated under CF condition exhibited more chalkiness and less transparency than those cultivated under GM condition. Scale bar, 10 mm. Percentage of (e) each category of grains and (f) protein contents in grains derived from rice plants cultivated in 2024 and 2025. Bar plots indicate mean values ± SD (*n* = 4 to 5). Asterisks and values above bars indicate significant differences (*P* < 0.05, Student’s *t*-test) and *P* values, respectively.

### Effects of hairy vetch treatment on nitrogen accumulation and leaf chlorophyll at panicle formation stage

Whole-shoot N concentration and flag leaf SPAD measured at the panicle formation stage, especially panicle initiation (around 40–50 DAT), serve as indicators of source capacity in rice (Yoshida, 1981; Ghosh et al., 2020). In our trials, both indicators were significantly higher with green manure than with chemical fertilizer (Fig. 3a), indicating more N available for producing grain storage proteins.

**Fig. 3.**
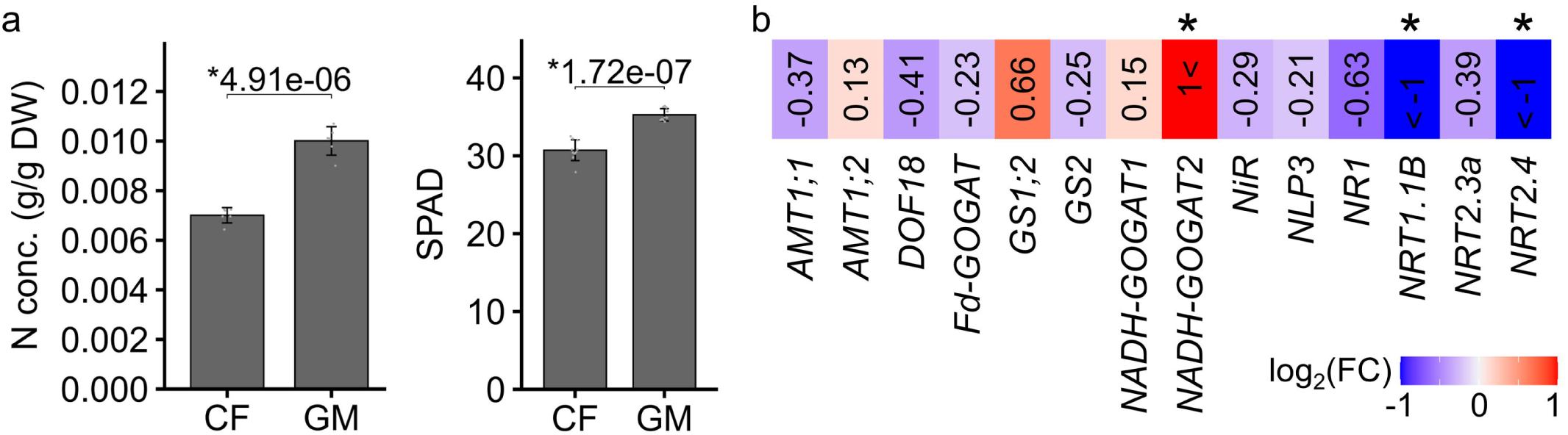
Nitrogen content indicators in shoots and nitrogen-related transcriptomes at a panicle formation stage. (a) Effects of GM treatment on total shoot nitrogen concentration and flag leaf SPAD values at panicle formation stage. Bar plots indicate mean values ± SD (*n* = 6). Asterisks and values above bars indicate significant differences (*P* < 0.05, Student’s *t*-test) and *P* values, respectively. (b) Changes in expression levels of well-known N-related genes detected in this study (raw read count is at least one in all the library samples). Asterisks represent genes showing statistically significant differences (|log_2_(FC)| > 1 and padj < 0.05).

### Effects of green manure treatment on leaf transcriptome profile at vegetative and panicle formation stages

To explore the genome-wide effect of green manure on transcriptome profiles, we performed RNA-seq using Koshihikari leaves at the vegetative and panicle formation stages and compared RNA-seq read counts for all well-expressed genes between treatments with chemical fertilizer and green manure. The principal component analysis suggested that the transcriptome profiles were changed in a fertilizer type–dependent manner (Supplementary Fig. 2a, Supplementary Fig. 3). With a threshold of more than two-fold change and padj < 0.05, 55 genes were significantly upregulated and 25 genes were significantly downregulated under green manure treatment compared to chemical fertilizer treatment in both 2023 and 2024 at the vegetative stage (Supplementary Fig. 2b). Enrichment GO analysis of the DEG set revealed that genes related to amino acid biosynthesis and photosynthesis were affected by fertilizer treatment type at the vegetative stage (Supplementary Fig. 2c). Among the genes corresponding to the enriched term ‘GO:0015979 photosynthesis’, several chloroplast precursors and chlorophyll biosynthetic gene *OsPORA* (Os04g0678700/LOC_Os04g58200) were significantly upregulated under green manure treatment (Supplementary Fig. 2d). Upregulated expression levels of these genes were consistent with the increased chlorophyll contents (Fig. 2a). We also investigated the changes in expression of previously established N-indicators that are upregulated under higher-N conditions (Takehisa and Sato, 2019). Transcripts of 7 out of 10 N-indicators exhibited significant increases under green manure treatment in both 2023 and 2024 (Supplementary Fig. 2e). Although the increases were not significant for the other three N-indicators in 2023, all were ∼2-fold or higher in both years. Furthermore, several well-known fertilizer-responsive genes or N-related genes showed significant transcriptional changes under the green manure treatment. For example, the *MORE AXILLARY GROWTH1-like* gene *Os1900* (Os02g0221900/LOC_Os02g45450), which participates in strigolactone biosynthesis and restricts tillering (Umehara et al., 2010; Cui et al., 2023), was significantly downregulated (Supplementary Fig. 2f). This tendency was consistent with increased tiller/panicle numbers (Fig. 2c). Notably, *OsNADH-GOGAT2* (Os05g0555600/LOC_Os05g48200), which positively influences N remobilization and loading into developing grains (Tamura et al., 2011), was significantly upregulated at both the vegetative and panicle formation stages (Fig. 3b, Supplementary Fig. 2g).

### Green manure–induced upregulation of *GA20ox2* and severe lodging

Despite its growth and yield advantages, the green manure treatment caused severe lodging in Koshihikari (Fig. 4a) that was associated with increased culm length (Fig. 4b), as reported in previous crop studies (Watanabe, 1984; Shah et al., 2017). In rice, there are at least four *GA20ox* genes, all positively affecting culm elongation (Oikawa et al., 2004; Asano et al., 2007; Qin et al., 2013; Wu et al., 2016; Agata et al., 2020). We explored whether expression levels of *GA20ox* genes were affected by green manure treatment. We detected *GA20ox1*, *GA20ox2*, and *GA20ox4* transcripts in all library samples (with a raw read count of at least one) (Fig. 4c). The expression level of only *Sd1*/*GA20ox2* was significantly upregulated under green manure treatment in 2023 (padj < 0.05, log_2_(FC) > 1). Although the increase in 2024 was not statistically significant, it was nearly 2-fold (log_2_(FC) = 0.82). *GA20ox1* and *GA20ox4* did not exhibit significant changes in either of the two years.

**Fig. 4.**
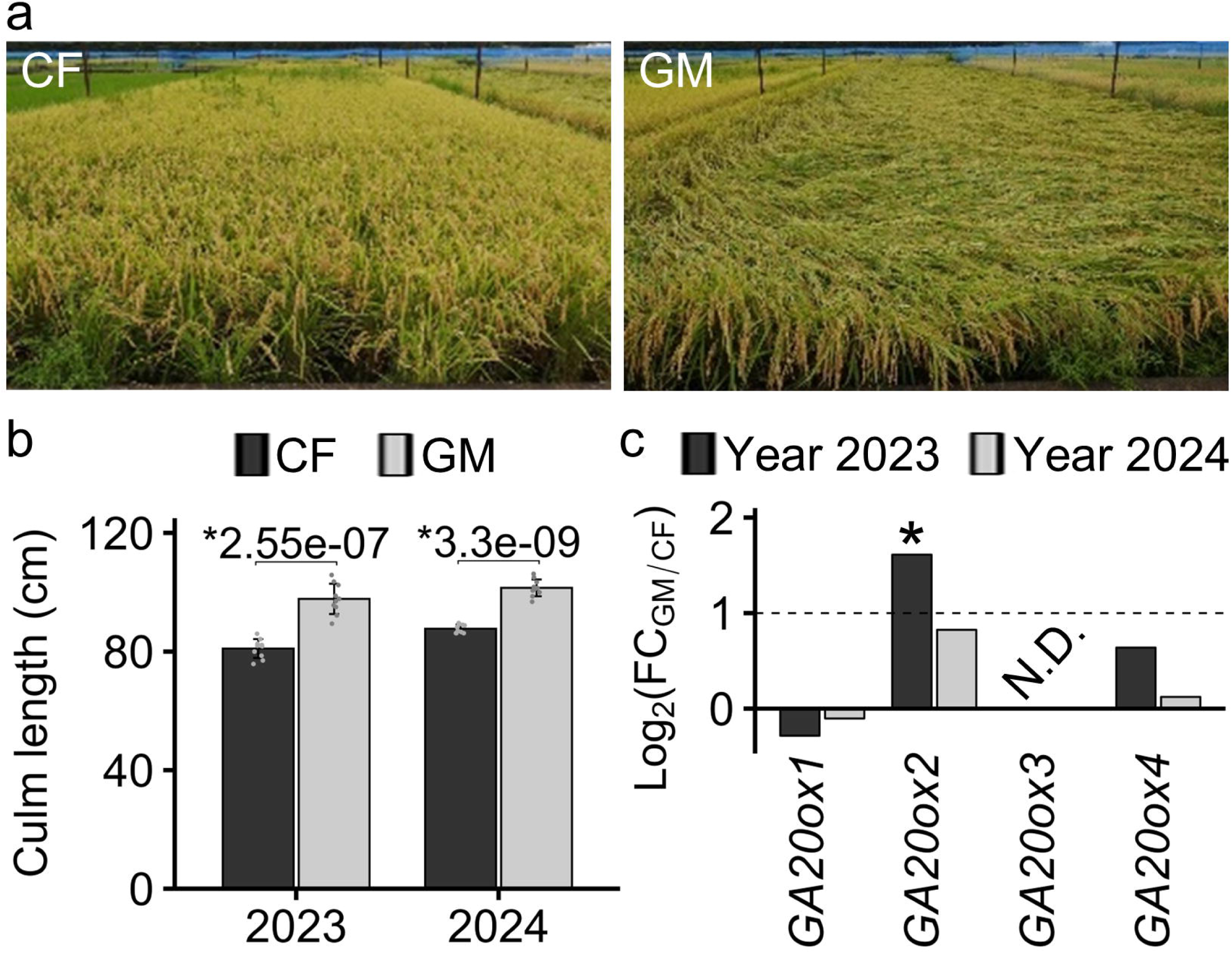
Severe effects of hairy vetch treatments on lodging in Koshihikari cultivated under field conditions. (a) Rice plants at 110 days after transplanting under chemical fertilization (CF) and green manure treatment (GM) conditions in 2023. Severe lodging was observed only under GM. (b) Culm length at harvesting stage in 2023 (1^st^ treatment) and 2024 (2^nd^ treatment; same treatment was applied to the plot treated in 2023). Bar plots indicate mean values ± SD (*n* = 10). Asterisks and numbers above each bar indicate significant differences (*P* < 0.05, Student’s *t*-test) and *P* values, respectively. (c) Changes in expression levels of *gibberellin 20 oxidase* (*GA20ox*) genes between CF and GM conditions. Asterisks represent DEGs (|log_2_(FC)| > 1 and padj < 0.05). N.D. represents less than 1 raw read count within any sample.

### Genotyping of *GA20ox2* to identify cultivars with possible lodging tolerance under green manure treatment

Since *GA20ox2* was upregulated in response to green manure treatment (Fig. 4c), we next analyzed the *GA20ox2* genotype in the same cultivars using PCR with previously established primers that detect a non-functional allele (∼380-bp deletion in the first exon) used to breed semi-dwarf Japanese cultivars (Asano et al., 2007). The results were compared between Koshihikari, which carries the non-dwarf (functional) allele (Tomita, 2009), and the other high-yielding Japanese cultivars including *indica* subgroup. The PCR products of Nijinokirameki, a temperate *japonica* cultivar, and Hokuriku 193, an *indica* cultivar, were about 360 bp (i.e., indicative of the deletion), while that of Koshihikari were 735 bp (Fig. 5a, b). This indicates that the alleles present in Nijinokirameki and Hokuriku 193 are of the non-functional type.

**Fig. 5.**
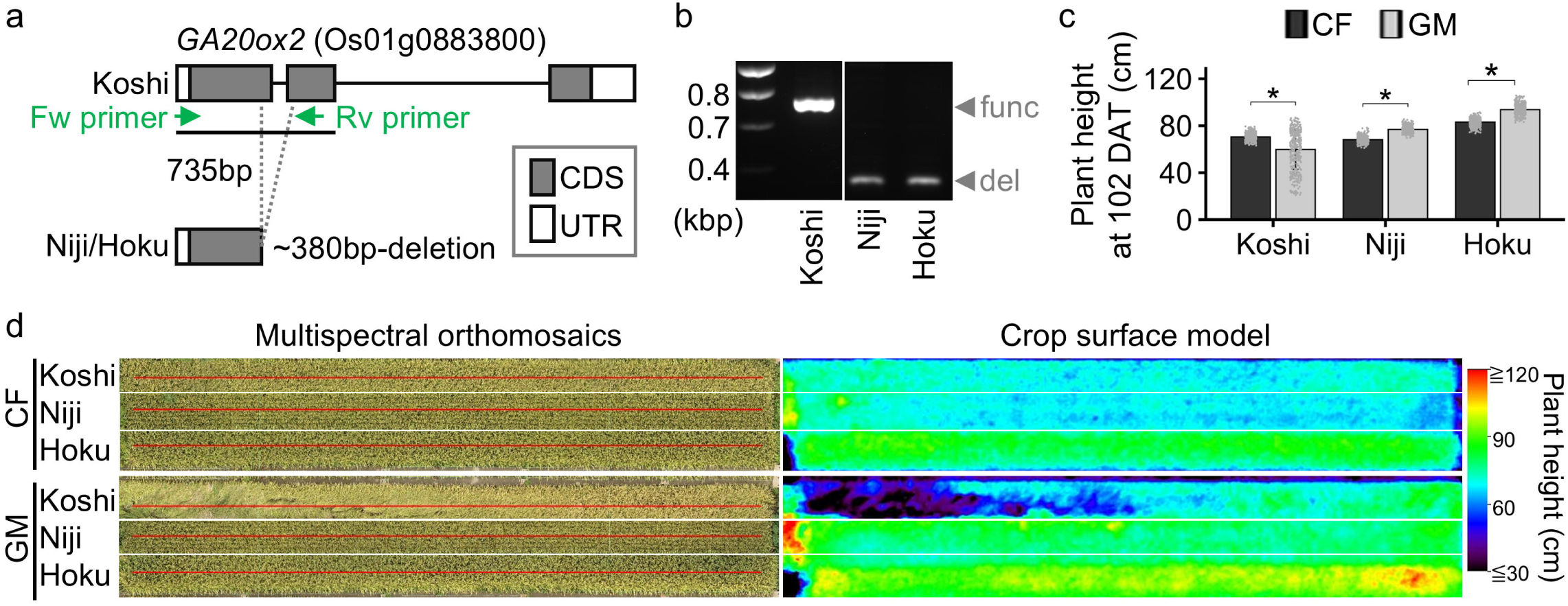
Lodging resistance in cultivars possessing non-functional allele of *GA20ox2* under green manure treatment. (a) Illustration of functional and non-functional alleles of *Sd1*/*GA20ox2*. Primer positions that facilitate the differentiation of these alleles are shown in green. (b) Genotyping of *gibberellin 20 oxidase 2* (*GA20ox2*) in leaves of different cultivars. func, 735-bp band indicating functional *GA20ox2* allele; del, ∼380-bp band indicating a non-functional *GA20ox2* deletion mutant allele. (c) Plant height of cultivars under field conditions at 102 days after transplanting (DAT) in 2024. The plant height was determined by UAV-based imaging. Two independent trials were performed for this investigation in different fields in the same period. The means ± SD are shown (*n* = 440). Asterisks indicate significant differences between chemical fertilization (CF) and green manure treatment (GM) in each cultivar (*P* < 0.05, Student’s *t*-test). (d) Multispectral orthomosaics and a crop surface model derived from aerial images obtained from UAV under CF and GM. The plants in the images are the same as those of (c). Red lines indicate the transects we used for the calculation of plant height shown in (c). White lines separate the different cultivars.

### Increased yields with lodging resistance in high-yielding Japanese cultivars harboring a non-functional *sd1*/*ga20ox2* allele under green manure treatment

Next, we compared lodging resistance in Koshihikari, Nijinokirameki, and Hokuriku 193 between the chemical fertilizer treatment and the green manure treatment. Plant heights calculated from aerial images were significantly decreased by green manure treatment in Koshihikari, but were significantly increased in the other two cultivars, at 102 DAT across two trials in 2024 (Fig. 5c). Nijinokirameki and Hokuriku 193 showed no lodging under either cultivation condition, while Koshihikari showed considerable lodging under the green manure treatment (Fig. 4a, Fig. 5d). Estimated grain yield (t/ha) of Nijinokirameki, and Hokuriku 193 were significantly higher under the green manure treatment than under the chemical fertilizer treatment, as were those of Koshihikari, in both 2023 and 2024 (Table 2).

**Table 2.** Estimated grain yield (t/ha) in different cultivars grown under CF and GM conditions in 2023 and 2024. The means ± SD are shown (*n* = 6, 2023; *n* = 5, 2024). Asterisks indicate significant differences between CF and GM conditions (*P* < 0.05, Student’s t-test).

| Grain yield (t/ha) |  |  |  |  |
| --- | --- | --- | --- | --- |
| Year | Condition | Koshi | Niji | Hoku |
| 2023 | CF | 4.7 ± 0.3 | 4.8 ± 0.4 | 7.2 ± 0.6 |
|  | GM (1<sup>st</sup>) | 7.4 ± 0.3* | 8.7 ± 0.5* | 9.9 ± 1.4* |
| 2024 | CF | 5.6 ± 0.6 | 5.5 ± 0.7 | 6.1 ± 0.9 |
|  | GM (2<sup>nd</sup>) | 7.4 ± 1.1* | 7.3 ± 0.3* | 8.9 ± 0.6* |

## Discussion

### Physiological effects of hairy vetch–based green manure treatment on vegetative growth and grain yield of rice under field cultivation

The application of green manure using leguminous cover crops such as hairy vetch is an environmentally friendly practice that reduces reliance on chemical fertilizers in rice cultivation; this is important because continuous use of chemical fertilizers reduces soil fertility (Hwang et al., 2015). In this study, we showed that hairy vetch–based green manure treatment promotes vegetative growth, and tiller number, at a level comparable to or even better than that obtained with chemical fertilizer treatment (Fig. 2a, c). The upregulation of high-N-responsive genes under green manure treatment (Fig. 3b, Supplementary Fig. 2e, g) suggests that N input exceeding 56 kg/ha was successfully obtained from hairy vetch plowed into field soils (Table 1). Additionally, leaf transcriptomic analysis revealed upregulation of genes related to chloroplast precursors and chlorophyll biosynthesis, resulting in increased total leaf chlorophyll content per unit area during the vegetative stage (Fig. 2a, Supplementary Fig. 2d). In general, higher chlorophyll content reflects better plant health during the growth stage (reviewed by Palta, 1990). These findings strongly suggest that hairy vetch applications can serve as an alternative to chemical fertilization for rice vigor promotion by increasing environmental N availability and supporting chlorophyll biosynthesis.

Next, we considered possible mechanisms other than chlorophyll biosynthesis by which hairy vetch–based green manure could increase grain yield. Our transcriptome analysis showed downregulated expression of *Os1900* under green manure relative to chemical fertilization (Supplementary Fig. 2f). Because *Os1900* negatively regulates tillering under nutrient limitation (Cui et al., 2023), its downregulation under a nutrient-replete regime would plausibly increase tiller/panicle number, enlarging sink capacity. We also observed upregulation of *OsNADH-GOGAT2*, a gene induced in expanded leaves at high N and associated with greater spikelet number and yield (Tamura et al., 2011), at the panicle formation stage (Fig. 3b). Tamura et al. (2011) inferred a role for OsNADH-GOGAT2 in converting leaf N into transportable glutamine during natural senescence, thereby enhancing phloem delivery of N to panicles and supporting grain filling. Consistent with this proposed mechanism, green manure increased whole-shoot N concentration and flag leaf SPAD at the panicle formation stage (Fig. 3a), indicating improved pre-senescence N status and a higher potential for N remobilization. Under that treatment, grain yield per plant increased without penalties of panicle length, grain size, and grain appearance quality (Fig. 2c–e, Supplementary Table 4), suggesting that yield increase was driven primarily by increased panicle number and, consequently, total spikelet number rather than by trade-offs that reduced resource allocation to grain production. Accumulating evidence may suggest a mechanism in which green manure increases sink capacity via *Os1900* downregulation and strengthens N recycling to developing grains via *OsNADH-GOGAT2* upregulation, leading to higher yields.

### Comparable grain yields achievements using hairy vetch–based green manure versus chemical fertilization under similar climate conditions

In our study, the maximum estimated grain yield was 9.9±1.4 t/ha in 2023 and 8.9±0.6 t/ha in 2024 (mean of two years: 9.4 t/ha), obtained through the cultivation of a recent high-yielding cultivar, Hokuriku 193 (Table 2). In a previous study based on chemical fertilizer use (80:28:40 kg/ha of N:PO_4_:K) performed in the same city (Tsukuba), Hokuriku 193 and its *MP3*-introgressed near-isogenic line exhibited estimated grain yields of 8.7 t/ha and 9.3 t/ha, respectively (Takai et al., 2024). This comparison suggests that hairy vetch–based green manure can achieve high yield levels more than those obtained through recent genetic improvement under conventional fertilizer-based management.

### A streamlined approach for selecting lodging-resistant rice cultivars under green manure treatment: focus on non-functional *sd1*/*ga20ox2* allele

The semi-dwarfing gene *sd1/ga20ox2* has been a cornerstone of modern rice breeding, particularly in Asia, where its loss-of-function alleles contributed to the development of high-yielding, lodging-resistant cultivars adapted to intensive fertilizer management (Ashikari et al., 2002; Asano et al., 2007; Sha et al., 2022). In the present study, *Sd1/GA20ox2* expression was upregulated under hairy vetch–based green manure treatment, suggesting that this GA biosynthetic gene responds to the nutrient-rich conditions created by green manure treatment. Because *Sd1*/*GA20ox2* is involved in GA biosynthesis and culm elongation, this transcriptional response may be associated with the increased lodging risk observed in cultivars carrying functional *Sd1*/*GA20ox2* alleles. In contrast, cultivars carrying a non-functional *sd1*/*ga20ox2* allele showed strong lodging resistance under hairy vetch–based green manure, indicating that this allele may contribute to stable plant architecture under green manure management. However, the physiological basis of lodging resistance under hairy vetch–based green manure remains to be clarified. Lodging-related traits such as pushing resistance, breaking resistance, culm morphology, and center-of-gravity height were not comprehensively evaluated in relation to *sd1*/*ga20ox2* function in this study and should be examined in future work. Overall, while further physiological validation is required, our results indicate that selecting cultivars carrying a non-functional *sd1*/*ga20ox2* allele is a practical approach for reducing lodging risk and stabilizing yield improvement even under hairy vetch–based green manure. These findings suggest that this well-established semi-dwarfing allele can be repurposed as a genetic resource for sustainable rice production systems that rely less on chemical fertilizer inputs.

## Conclusion

This study demonstrates that hairy vetch–based green manure treatment can enhance rice productivity, but that its benefits depend strongly on the genetic control of lodging resistance. Under green manure treatment, several high-yielding Japanese cultivars carrying a non-functional *sd1/ga20ox2* allele achieved estimated grain yields exceeding 9 t ha⁻¹ without severe lodging, reaching levels comparable to those obtained under chemical fertilizer management. Green manure treatment likely contributed to yield improvement by enhancing nitrogen availability during reproductive development, as supported by the upregulation of *OsNADH-GOGAT2* at the panicle formation stage. These findings indicate that the yield-promoting and quality-improving effects of green manure can be effectively captured when excessive culm elongation is suppressed by an appropriate semi-dwarf genetic background. Thus, selection of cultivars carrying non-functional *sd1/ga20ox2* alleles provides a practical route to overcome the major trade-off of green manure management: enhanced vegetative growth versus increased lodging risk. This study will facilitate the implementation of annual hairy vetch–rice rotation systems for high-yield rice production with reduced reliance on chemical fertilizers.

## Abbreviations

CI_green_: Chlorophyll Index-Green
CF: Chemical fertilizer
CSM: crop surface model
DNA: Deoxyribonucleic acid
DAT: days after transplanting
DSM: digital surface model
GA: Gibberellin
GCP: ground control point
GM: Green manure
GO: Gene ontology
K: Pottassium
N: Nitrogen
NIR: Near-Infrared
PO_4_: Phosphate ion
RNA: Ribonucleic acid
RNA-seq: RNA-sequencing
SfM-MVS: Structure-from-Motion/Multi-View-Stereo
SPAD: Soil Plant Analysis Development

## Declarations

### Ethics approval and consent to participate

Not applicable

### Consent for publication

Not applicable

### Availability of data and materials

All data generated or analyzed during this study are included in this published article and its supplementary information files.

### Authorship contribution

**Hirofumi Fukuda:** Writing – original draft, Writing – review & editing, Investigation, Data curation, Formal analysis, Visualization, Conceptualization. **Toshihiro Sakamoto:** Writing – review & editing, Investigation, Data curation, Formal analysis. **Akari Fukuda:** Investigation, Data curation. **Daisuke Ogawa:** Writing – review & editing, Investigation, Data curation, Funding acquisition, Conceptualization, Supervision.

## Acknowledgements

We thank Makiko Suzuki, Yuko Aono, and Aimi Murakami for their technical and field support. We would like to thank Kazuho Funakawa for helping us analyze drone data. We appreciate the technical staff of the Institute of Crop Science, NARO, for their help in managing the rice fields. We would like to thank Dr. Kawakatsu for preparing the RNA sequence library. We also thank Kett Electric Laboratory Co. Ltd. for helping with the analysis of protein content of rice grains. We thank Snow Brand Seed Co., Ltd. for helping with hairy vetch cultivation. We are thankful to the editors at ELSS, Inc. (https://elss.co.jp/en/) for their professional editing services before submission.

## Declaration of generative AI and AI-assisted technologies in the writing process

During the preparation of this work, we used ‘DeepL Write’, ‘Microsoft Copilot’ and ‘ChatGPT’ in order to improve the grammar, spelling, and readability. After using these tools, the authors carefully reviewed and edited the content as needed and take full responsibility for the content of the published article.

