## Supplementary_Figs for "Rice Cultivars Carrying the Semi-Dwarfing Allele Enables High Yield without Lodging under Hairy Vetch–based Green Manure"

### Slide 1
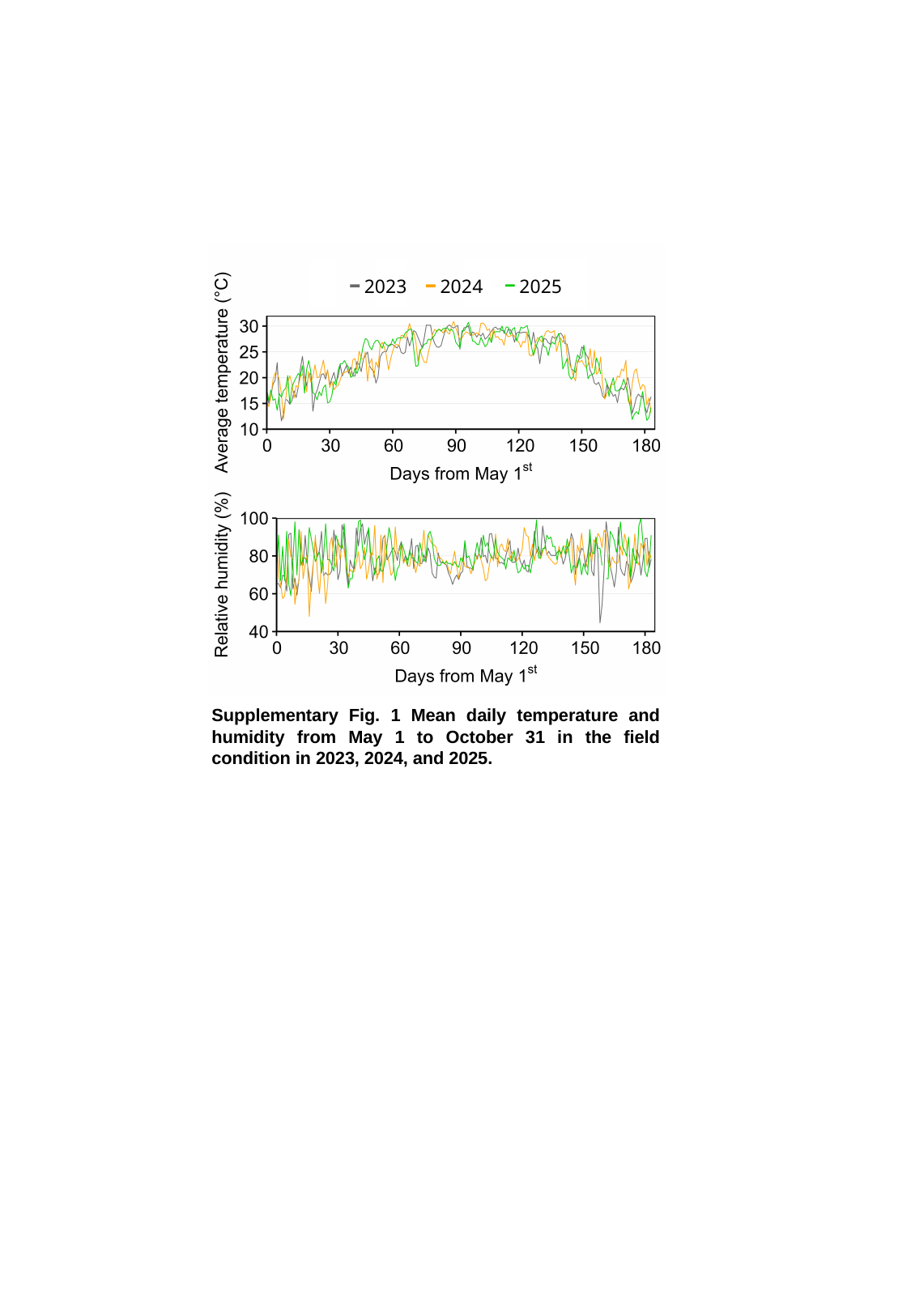

2023
2024
2025
Supplementary Fig. 1 Mean daily temperature and humidity from May 1 to October 31 in the field condition in 2023, 2024, and 2025.

### Slide 2
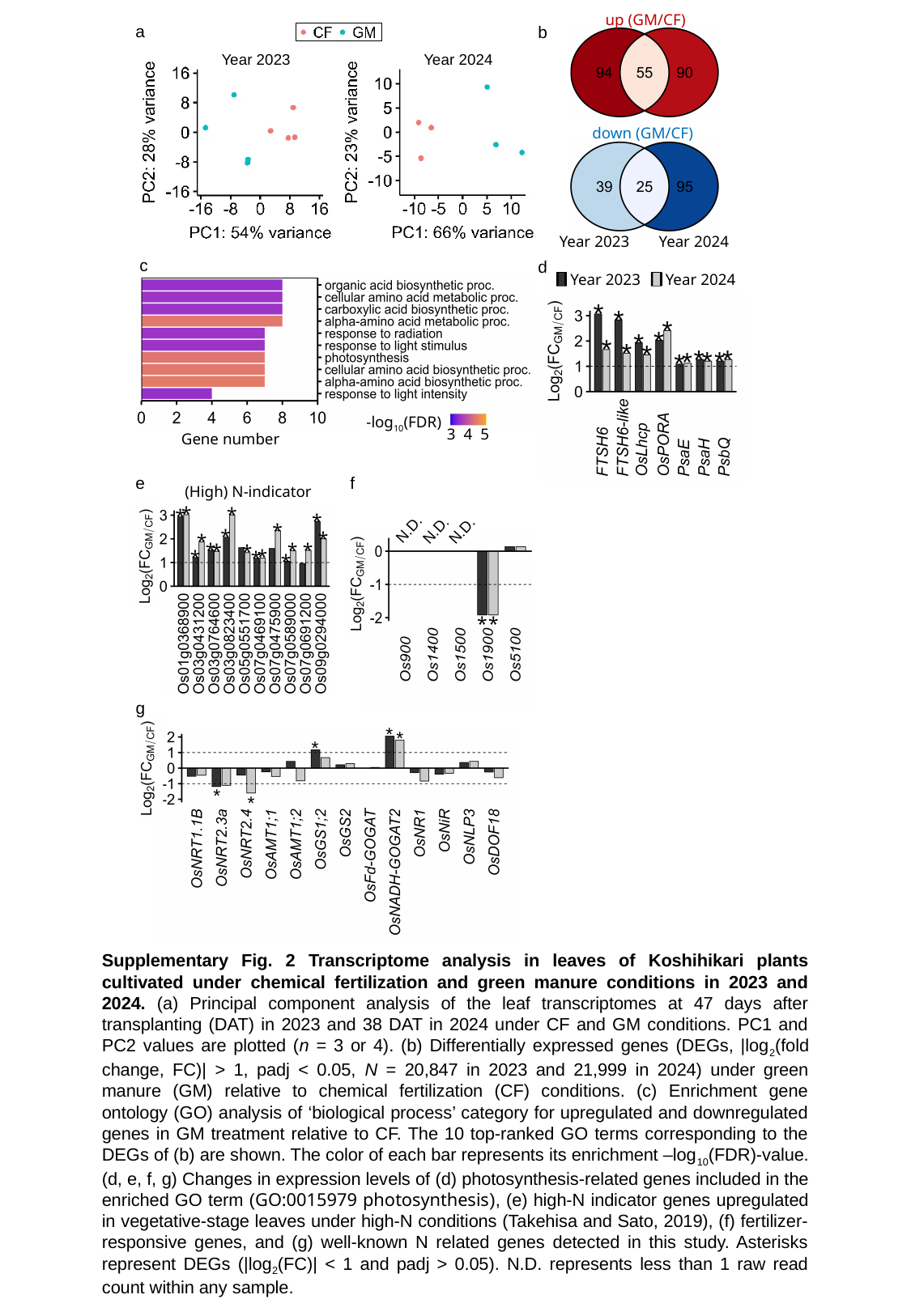

up (GM/CF)
a
b
Year 2023
Year 2024
down (GM/CF)
Year 2024
Year 2023
c
d
Year 2023
Year 2024
*
*
*
*
*
*
*
*
*
*
*
*
*
*
-log_10_(FDR)
3 4 5
Gene number
e
f
(High) N-indicator
*
*
*
*
*
N.D.
N.D.
N.D.
*
*
*
*
*
*
*
*
*
*
*
*
*
*
g
*
*
*
*
*
Supplementary Fig. 2 Transcriptome analysis in leaves of Koshihikari plants cultivated under chemical fertilization and green manure conditions in 2023 and 2024. (a) Principal component analysis of the leaf transcriptomes at 47 days after transplanting (DAT) in 2023 and 38 DAT in 2024 under CF and GM conditions. PC1 and PC2 values are plotted (n = 3 or 4). (b) Differentially expressed genes (DEGs, |log_2_(fold change, FC)| > 1, padj < 0.05, N = 20,847 in 2023 and 21,999 in 2024) under green manure (GM) relative to chemical fertilization (CF) conditions. (c) Enrichment gene ontology (GO) analysis of ‘biological process’ category for upregulated and downregulated genes in GM treatment relative to CF. The 10 top-ranked GO terms corresponding to the DEGs of (b) are shown. The color of each bar represents its enrichment –log_10_(FDR)-value. (d, e, f, g) Changes in expression levels of (d) photosynthesis-related genes included in the enriched GO term (GO:0015979 photosynthesis), (e) high-N indicator genes upregulated in vegetative-stage leaves under high-N conditions (Takehisa and Sato, 2019), (f) fertilizer-responsive genes, and (g) well-known N related genes detected in this study. Asterisks represent DEGs (|log_2_(FC)| < 1 and padj > 0.05). N.D. represents less than 1 raw read count within any sample.

### Slide 3
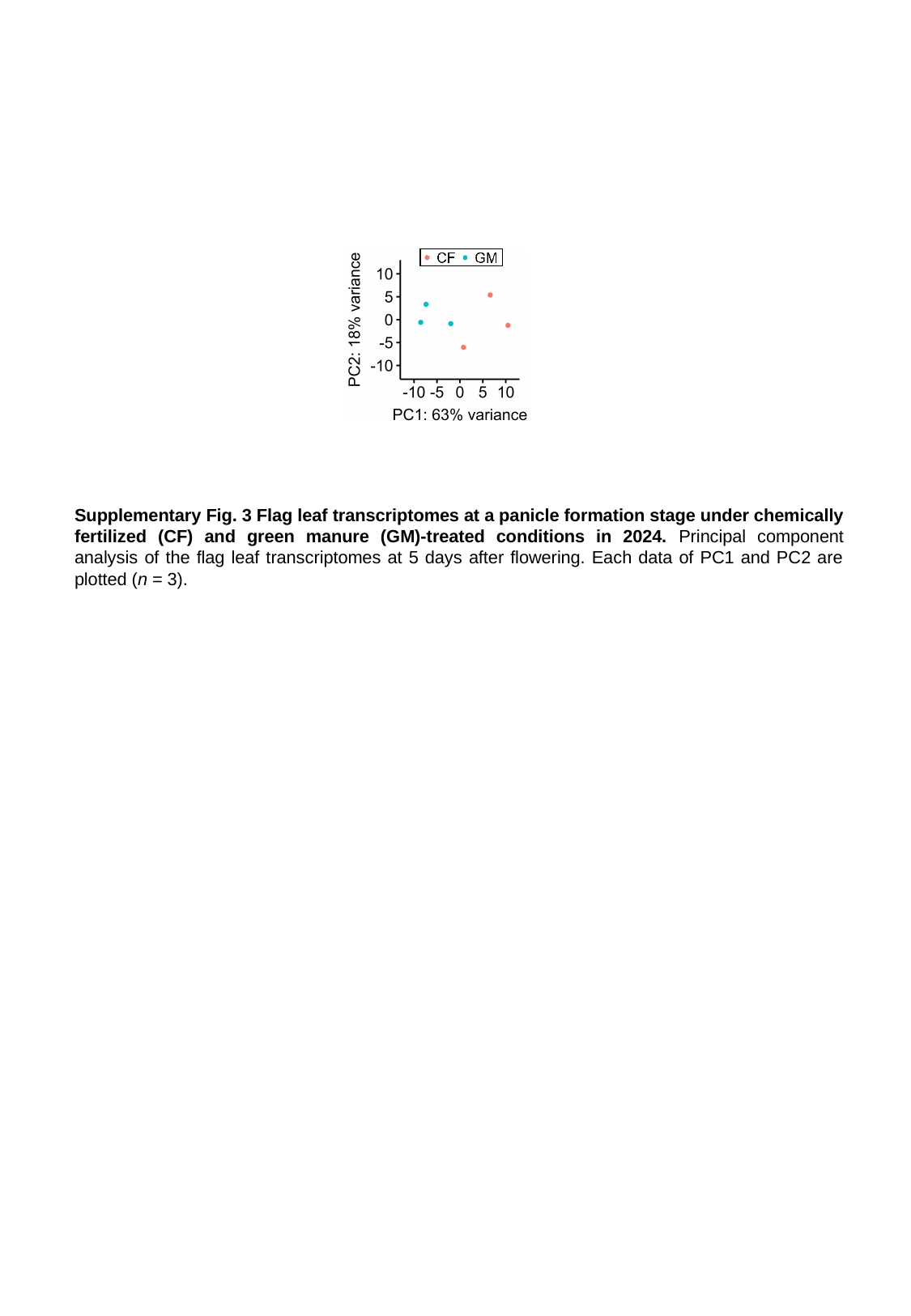

Supplementary Fig. 3 Flag leaf transcriptomes at a panicle formation stage under chemically fertilized (CF) and green manure (GM)-treated conditions in 2024. Principal component analysis of the flag leaf transcriptomes at 5 days after flowering. Each data of PC1 and PC2 are plotted (n = 3).
